# Broad-spectrum antivirals of protoporphyrins inhibit the entry of highly pathogenic emerging viruses

**DOI:** 10.1101/2020.05.09.085811

**Authors:** Shengsheng Lu, Xiaoyan Pan, Daiwei Chen, Xi Xie, Yan Wu, Weijuan Shang, Xiaming Jiang, Yuan Sun, Sheng Fan, Jian He

**Affiliations:** Group of Peptides and Natural Products Research, School of Pharmaceutical Sciences, Southern Medical University, 1838 Guangzhou Avenue North; Guangzhou 510515, China; State Key Laboratory of Virology, Wuhan Institute of Virology, Center for Biosafety Mega-Science, Chinese Academy of Sciences, Wuhan 430071, China; University of the Chinese Academy of Sciences, Beijing, China

**Keywords:** Protoporphyrin IX (**PPIX**), Lipid biolayer-targeting, Broad-spectrum antivirals, Enveloped virus

## Abstract

Severe emerging and re-emerging viral infections such as Lassa fever, Avian influenza (AI), and COVID-19 caused by SARS-CoV-2 urgently call for new strategies for the development of broad-spectrum antivirals targeting conserved components in the virus life cycle. Viral lipids are essential components, and viral-cell membrane fusion is the required entry step for most unrelated enveloped viruses. In this paper, we identified a porphyrin derivative of protoporphyrin IX (**PPIX**) that showed broad antiviral activities *in vitro* against a panel of enveloped pathogenic viruses including Lassa virus (LASV), Machupo virus (MACV), and SARS-CoV-2 as well as various subtypes of influenza A viral strains with IC_50_ values ranging from 0.91±0.25 μM to 1.88±0.34 μM. A mechanistic study using influenza A/Puerto Rico/8/34 (H1N1) as a testing strain showed that **PPIX** inhibits the infection in the early stage of virus entry through biophysically interacting with the hydrophobic lipids of enveloped virions, thereby inhibiting the formation of the negative curvature required for fusion and blocking the entry of enveloped viruses into host cells. In addition, the preliminary antiviral activities of **PPIX** were further assessed by testing mice infected with the influenza A/Puerto Rico/8/34 (H1N1) virus. The results showed that compared with the control group without drug treatment, the survival rate and mean survival time of the mice treated with **PPIX** were apparently prolonged. These data encourage us to conduct further investigations using **PPIX** as a lead compound for the rational design of lipid-targeting antivirals for the treatment of infection with enveloped viruses.

## Introduction

Outbreaks of severe pathogenic viral infections such as Avian influenza (AI), severe acute respiratory syndrome-coronavirus, Middle East Respiratory Syndrome-coronavirus, and Ebola virus have posed a significant threat to public health. Among them, the outbreaks this year of Lassa fever in Nigeria caused by the deadly Lassa virus has infected 472 people and led to more than 70 deaths; influenza induced by influenza virus in the United States has infected more than 25 million people and caused 14,000 deaths. Pneumonia due to infection with the novel coronavirus SARS-CoV-2 (also named COVID-19), which is currently circulating globally, has caused more than 230,000 deaths. These outbreaks highlight the urgent need for new strategies and approaches to developing efficient antiviral drugs with broad-spectrum activities for prophylactic and therapeutic usage. Notably, many of these viruses are enveloped RNA viruses, and a third of the currently emerging and re-emerging infections are caused by this type of pathogen (1). As such, we speculated that it might be possible to develop an arsenal of broad-spectrum antivirals by targeting conserved components involved in the life cycle of these viruses.

Structurally, an enveloped virus particle comprises a genome and a protein coat or capsid that are surrounded by a lipid bilayer envelope, where the genome encoding viral elements is enclosed inside the capsid compartment (2,3). In the virus life cycle, the entry process is the first step, which involves multiple scenarios, including attachment to the surfaces of host cells, internalization into host cell compartments by exploitation of cellular uptake machineries (4), transport to the lumen of endosomes or the endoplasmic reticulum (ER) (5), and fusion of viral and host cell membranes (6). Due to their crucial roles in mediating the entry of a virus, each of these steps is a promising target for antivirals, which can intervene by blocking the infection in the early stage. Among these steps, the fusion of the virus envelope with the cell membrane is a known essential step for all enveloped viruses (7) and some molecules targeting this step have shown broad antiviral activity (8–10).

Porphyrins and their derivatives are a group of conjugated planar molecules that possess broad applications in the field of biomedical science, materials chemistry, and electrochemistry due to their unique spectroscopic properties and electrochemical performance (11–14). Of particular interest for biomedical science is their significant role in therapies that are associated with microbial infection and tumor therapies (15–16). In our previous work, through a bioassay-guided approach, we identified a porphyrin derivative of pyropheophorbide a (**PPa**) from the marine mussel *Musculus senhousei.* The biological activity test showed that **PPa** exhibited good antiviral activity against a panel of influenza A viruses. The mechanism of action study indicated that **PPa** exerted its inhibitory effect in the early stage of virus infection by interacting with the lipid bilayer function of the virion, blocking the entry of enveloped viruses into host cells (17). Considering the proposed drug target of **PPa**, we decided to conduct extensive research by synthesizing a number of derivatives possessing the same skeleton of porphyrin as that of **PPa** to study its broad antiviral activities in addition to those exhibited against influenza A viruses. Hence, an easily accessed material, protoporphyrin IX (**PPIX**) (Fig. 1A), was adopted as a starting material, and derivatives were designed by increasing the hydrophobicity of **PPIX** with the expectation of increasing the interactions between **PPIX** and the lipid of the virus. These molecules were then tested for antiviral activities against a broad panel of enveloped viruses including the highly pathogenic Lassa virus (LASV) and Machupo virus (MACV) as well as SARS-CoV-2 which causes COVID-19 in China currently. In addition, various subtypes of influenza A viruses (IAV) were also included. Herein, we report the antiviral activities as well as the possible modes of action of these molecules.

**Fig. 1.**
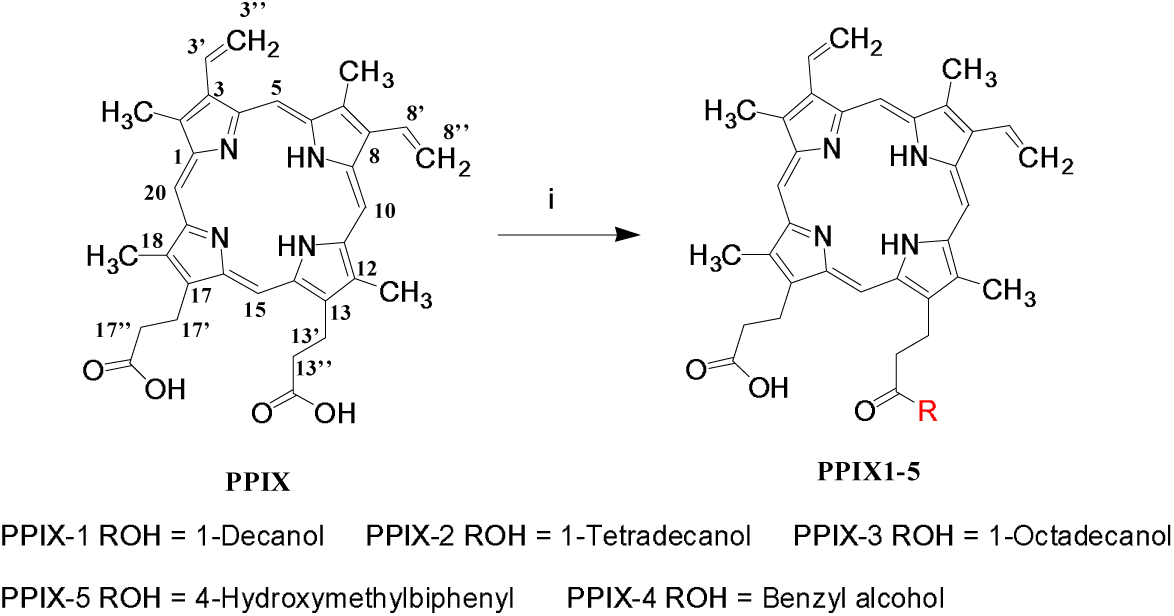
The structure of **PPIX** and its derivatives. Reagents and conditions: (i) DMF, HBTU, DMAP, DCC, DIEA, 0[for the first 30 min, then turned to room temperature.

## Results

### Protoporphyrin IX derivatives were synthesized by reacting with various alcohols

Protoporphyrin IX (**PPIX**) is a heterocyclic molecule consisting of four pyrrole rings and two symmetric carboxyl groups (Fig. 1), which were then used as a functional group to conjugate with other alcohols. Considering the possible target of **PPa** as the lipid bilayer of the virion, alcohols with various lengths of aliphatic chains or aromatic rings were used to react with **PPIX** to enhance the hydrophobicity and steric effects, thereby studying the antiviral activities of these compounds.

As a result, two types of **PPIX** derivatives (**1-5**) including three aliphatic and two aromatic alcohols were synthesized, as indicated in Fig. 1. The obtained derivatives were purified, their structures were confirmed by NMR and MS spectroscopic data, and they were used for the following antiviral activity evaluation.

### PPIX showed broad-spectrum antiviral activities against a panel of unrelated enveloped viruses

Compounds **1-5** together with **PPIX** were tested for their antiviral activity against a number of enveloped viruses including pseudotyped Old World and New World human arenaviruses Lassa virus (LASV) and Machupo virus (MACV), respectively, and various subtypes of influenza A viruses (IAV), such as A/Puerto Rico/8/34 (H1N1), A/FM/1/47 (H1N1) mouse-adapted viral strain, A/Puerto Rico/8/34 (H1N1) with the NA-H274Y mutation, and A/Aichi/2/68 (H3N2) viral strains, as indicated in Table 1. By using a ‘pretreatment of virus’ approach (18), the antiviral efficacies were assessed by measuring the survival of the host cells using MTT assay in an anti-IAV activity test and measuring the luciferase activity toward the pseudotyped LASV and MACV, as described before (19).

**Table 1.** Inhibitory activity of **PPIX** and its derivatives against various viruses and cellular toxicities

| Name | IC<sub>50</sub>±SD (μM)<sup>a</sup> |  |  |  |  |  | CC<sub>50</sub>±SD (μM)<sup>b</sup> |  | MW<sup>h</sup> |
| --- | --- | --- | --- | --- | --- | --- | --- | --- | --- |
|  | PR8<sup>c</sup> | FM-1<sup>d</sup> | 274<sup>e</sup> | H3N2<sup>f</sup> | LASV<sup>g</sup> | MACV<sup>g</sup> | MDCK | Vero-E6 |  |
| **PPIX** | 1.88±0.85 | 1.88±0.34 | 1.24±0.53 | 1.49±0.66 | 0.91±0.25 | 1.07±0.27 | 229.59±4.53 | 120.80±0.95 | 562.66 |
| **PPIX-1** | 7.2±3.23 | 12.12±2.68 | 13±1.36 | 8.95±1.68 | 4.63±2.16 | 17.34±3.88 | 121.23±2.18 | 76.39±2.29 | 720.94 |
| **PPIX-2** | 21.27±3.5 | >25.74 | 21.12±2.16 | 13.63±2.01 | >25.74 | >25.74 | 128.32±0.54 | 111.90±3.58 | 777.05 |
| **PPIX-3** | >21.93 | >21.93 | >21.93 | >21.93 | >21.93 | >21.93 | 139.56±1.58 | 127.970±1.46 | 912.16 |
| **PPIX-4** | >29.82 | >29.82 | >29.82 | >29.82 | >29.82 | >29.82 | 198.57±1.62 | 137.70±2.89 | 670.79 |
| **PPIX-5** | >26.78 | >26.78 | >26.78 | >26.78 | >26.78 | >26.78 | 237.49±2.72 | 215.93±2.18 | 746.89 |
| **PPa** | 0.32±0.28 | 1.65±0.56 | 1.05±0.99 | 1.23±1.10 | NT<sup>i</sup> | NT | 139.98±3.61 | NT | 534.66 |
| **S-KKWK<sup>i</sup>** | 1.85±0.92 | 1.96±0.23 | NT | NT | NT | NT | NT | NT | 1014.84 |

<sup>a</sup> The anti-influenza A virus (IAV) activity was determined after the virus was pretreated with **PPIX** or its derivatives.
<sup>b</sup> The toxicities of MDCK or Vero-E6 cells were assessed using MTT assay.
<sup>c</sup> A/Puerto Rico/8/34, <sup>d</sup> A/FM/1/47 mouse-adapted viral strain, <sup>e</sup> A/Puerto Rico/8/34 with the NA-H274Y mutation, <sup>f</sup> A/Aichi/2/68. <sup>g</sup> The activity was assessed by measuring the luciferase activity toward the pseudotyped LASV and MACV.
<sup>h</sup> MW: molecular weight, <sup>i</sup> NT: not tested.
<sup>i</sup> S-KKWK was used as a positive control, as reported previously (20).

The data in Table 1 showed that the prototype of **PPIX** was the most active one among all tested porphyrins, displaying broad activity against all viral strains with IC_50_ values ranging from 0.91 to 1.88 μM. Significantly, **PPIX** was also active toward the oseltamivir-resistant virus of influenza A/Puerto Rico/8/34 (H1N1) with the NA-H274Y mutation.

To confirm the antiviral effect of **PPIX**, we focused on influenza A virus and employed ‘pretreatment of the virus’ and ‘treatment during infection’ methods to test the inhibitory activity of **PPIX** using indirect immunofluorescence staining, qRT-PCR and plaque reduction assays. As shown in Fig. 2A, the data from the indirect immunofluorescence staining on the expression of the viral nucleoprotein (NP) in MDCK cells showed that the green fluorescence associated with NP expression from the A/Puerto Rico/8/34 (H1N1) influenza virus was apparently decreased after treatment with **PPIX**. Moreover, the ‘Pretreatment’ group showed more potent than the ‘During infection’ group, which was further qualified as indicated in Fig. 2B using ImageJ software (21).

**Fig. 2.**
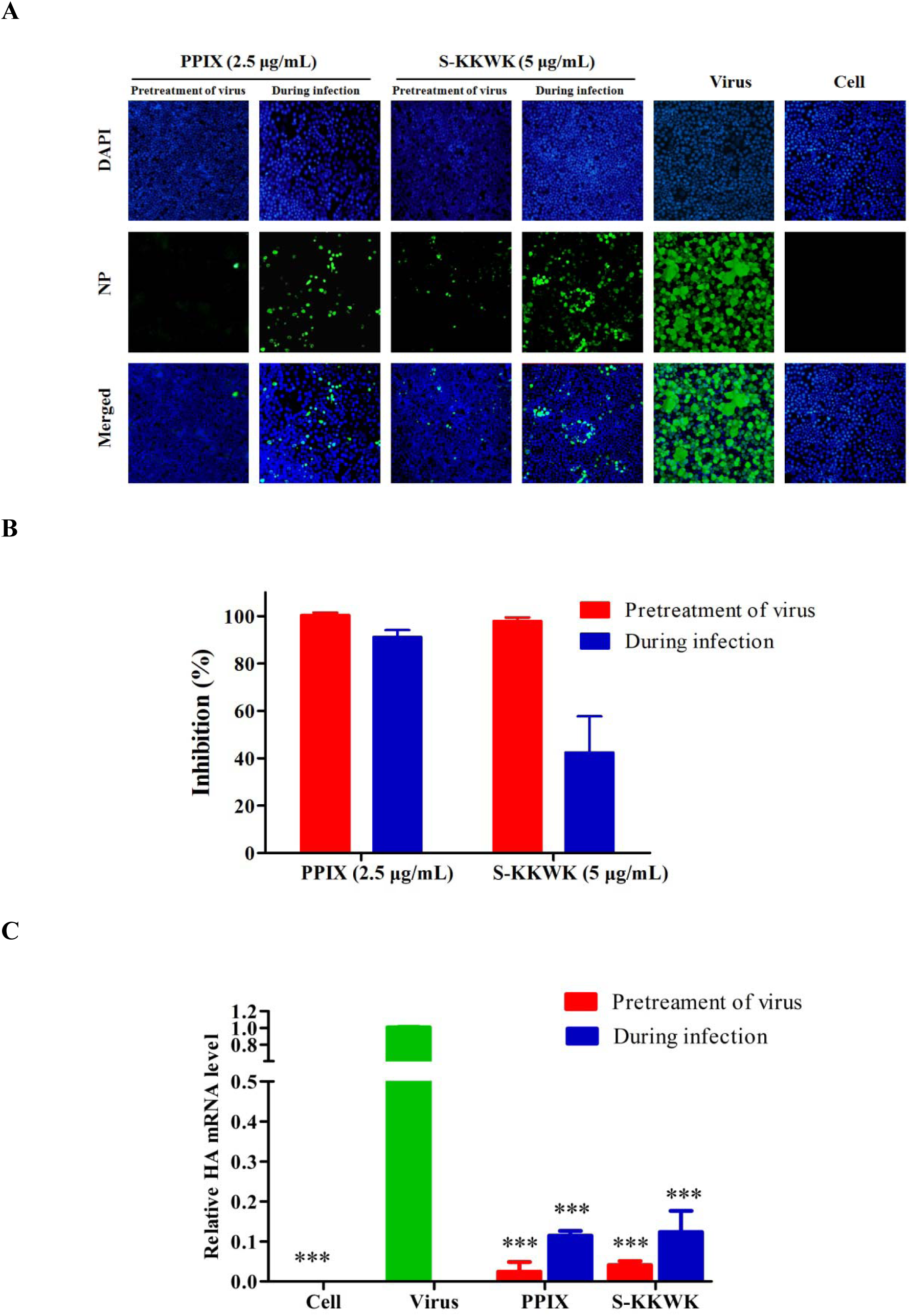
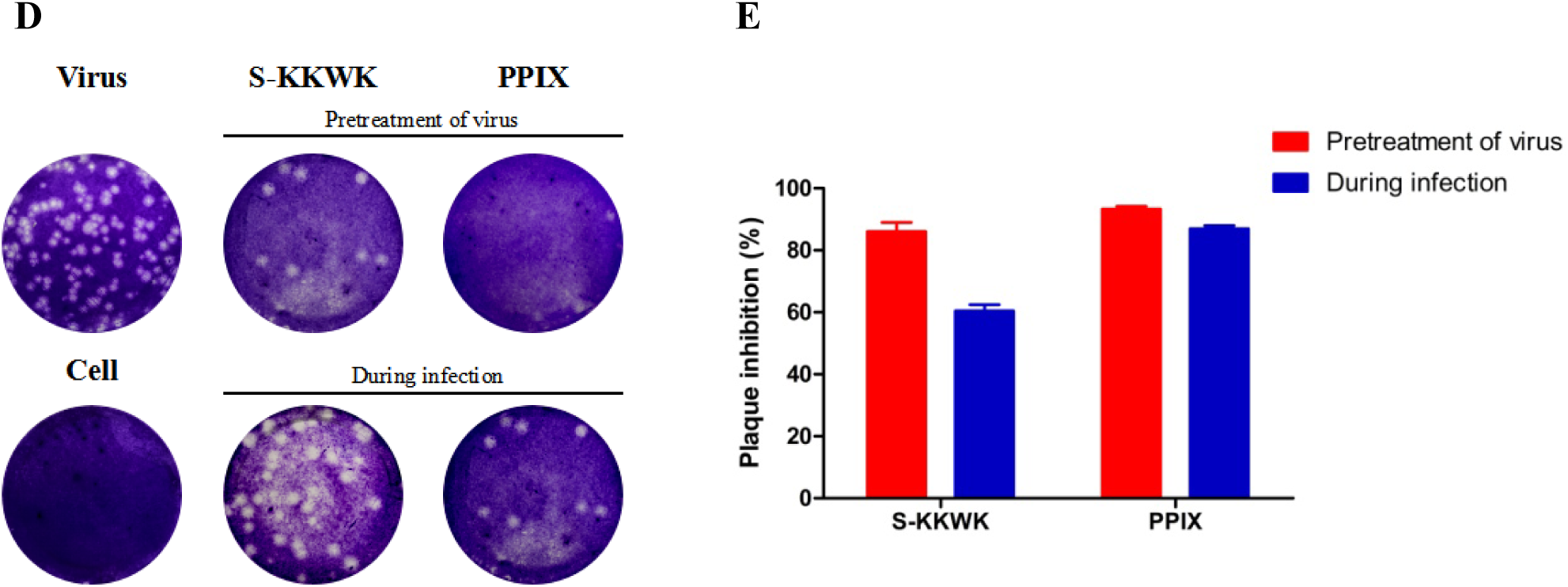
**A** The inhibitory effect of **PPIX** on the expression of NP in MDCK cells using indirect immunofluorescence assay. In this assay. 100 TCID_50_ of influenza A/PR/8/34(H1N1) virus and 2.5 μg/mL **PPIX** were used to treat MDCK cells by ‘pretreatment of virus’ and ‘during infection’ approaches, respectively. At 24 h p.i, the expression of NP (green fluorescence) in the cytoplasm of MDCK cells was immunostained and observed under a fluorescence microscope. S-KKWK (5 μg/mL) was used as a positive control. DAPI staining (blue fluorescence) was employed to indicate the position of the nucleus. **2B** The NP expression (green fluorescence) in each group was quantified using ImageJ software, and the inhibitory effect of **PPIX** on NP expression was thus calculated. **2C** The mRNA levels of the HA gene after treatment with **PPIX** using the ‘pretreatment of the virus’ or ‘during infection’ drug administration approaches were used to evaluate the antiviral effects of **PPIX** on influenza A/PR/8/34 (H1N1). The significance of the differences in the data between the virus-infected groups was determined using one-way ANOVA: ^*^ *p*< 0.05, ^**^ *p*< 0.01, *and* ^***^ *p*< 0.001. **2D** Plaque reduction assay to test the inhibitory effects of **PPIX** (5 μg/mL) to reduce the number of plaques of the influenza A/Puerto Rico/8/34 (H1N1) virus using the ‘pretreatment of virus’ and ‘during infection’ approaches. In the first approach, **PPIX** was preincubated with 100 TCID_50_ of the virus for 30 min at 37℃ before being added to the cells, whereas in the latter approach virus mixed with **PPIX** was added to the cells simultaneously. After 48 h p.i, the cells were fixed and stained. **2E** The number of plaques in each group were counted, from which the plaque inhibition rate of **PPIX** against the influenza A/Puerto Rico/8/34 (H1N1) virus was obtained.

Moreover, both measurements of the mRNA level of the HA gene from influenza A/PR/8/34 (H1N1) after treatment with **PPIX** (qRT-PCR) (Fig. 2C) and the plaque reduction assay (Fig. 2D and **2E**) showed similar results as the indirect immunofluorescence staining (Fig. 2A), which again confirmed the antiviral effects of **PPIX**.

Furthermore, to confirm the broad antiviral activity of **PPIX**, we tested its activity toward the recently emerged deadly virus Severe Acute Respiratory Syndrome Coronavirus 2 (SARS-CoV-2). Two drug administration approaches including ‘Full-time treatment’ and ‘Pretreatment of virus’ were employed to test the inhibitory effect of **PPIX** toward this virus. As shown in Fig. 3A, the obtained IC_50_ values were 0.11±0.02 μM and 0.21±0.02 μM, respectively, compared with 3.7 μM for Remdesivir and 10 μM for chloroquine under the same ‘Full-time treatment’ testing condition (22). Moreover, the *in vitro* anti-SARS-CoV-2 activity was again assessed by immunofluorescence microscopy toward this virus using the ‘Full-time treatment’ condition and detected at 48□h p.i (Fig. 3B).

**Fig. 3.**
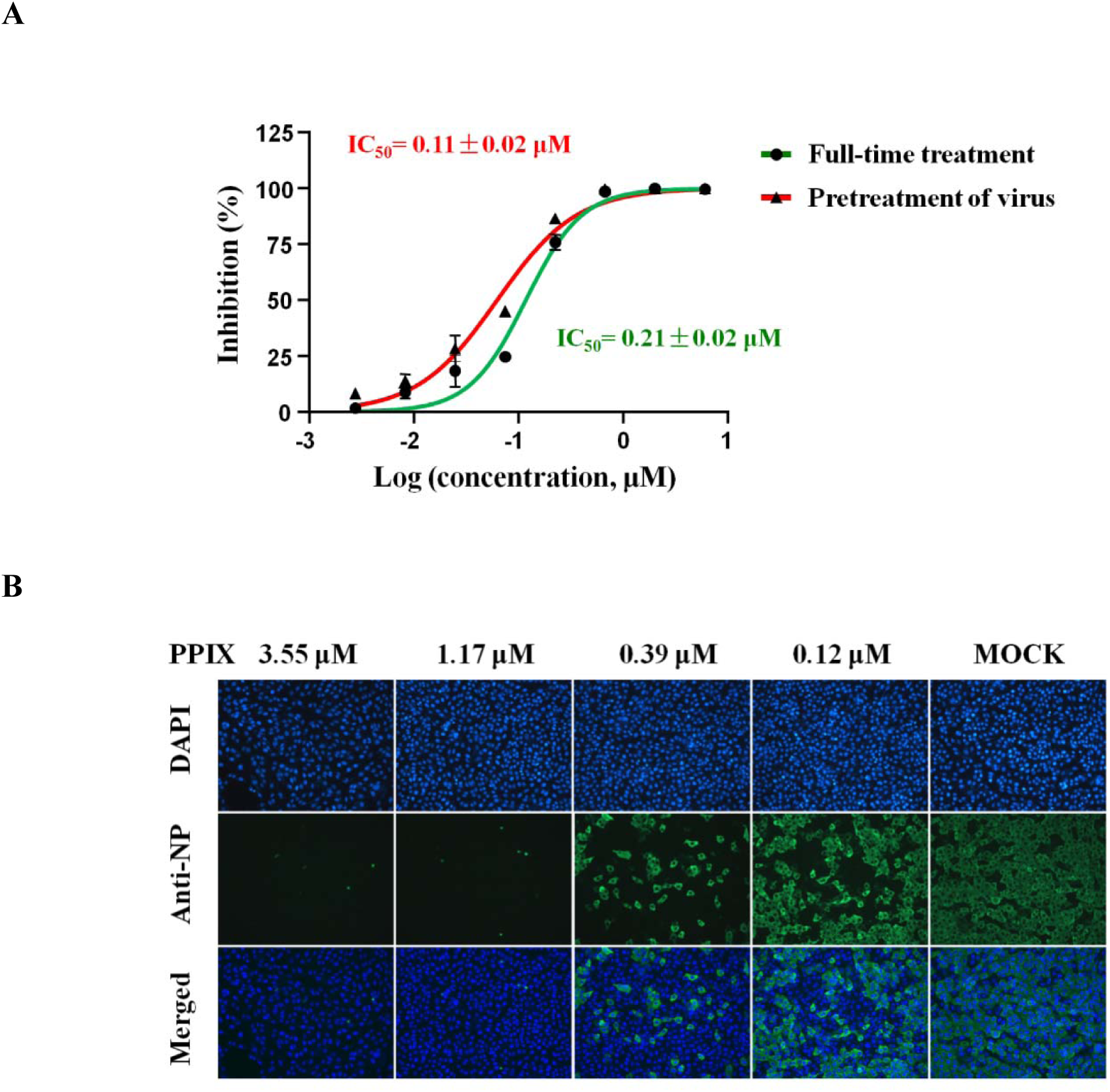
The inhibitory effect of **PPIX** against SARS-CoV-2. **3A** Two drug administration approaches including ‘Full-time treatment’ and ‘Pretreatment of virus’ were used in the experiment. In the first method, 5×10^4^ overnight cultured Vero E6 cells/well were pretreated with the serially diluted **PPIX** for 1 h, followed by the addition of virus (MOI of 0.05) and incubation for 2 h. After removal of the virus-**PPIX** mixture, the cells were cultured with fresh **PPIX**-containing medium for 24 h postinfection. In the ‘Pretreatment of virus’ approach, the serially diluted **PPIX** was preincubated with virus (MOI of 0.05) for 1 h at 37°C prior to being added into the cells. After infection for 1 h, the virus-**PPIX** mixture was removed and fresh medium was added to the cells, which were allowed to incubate at 37□ in a 5% CO_2_ atmosphere for 24 h. In both approaches, at 24 h p.i the cell supernatant was collected and the S gene of the virus was quantified with qPCR^21^; then, the inhibitory efficacies represented as IC_50_ values were calculated using MOCK as a blank control. **3B** The anti-SARS-CoV-2 activity on the expression of NP in Vero E6 cells using indirect immunofluorescence assay. Following the same full-time treatment protocol and after incubation of cells for 24 h of postinfection, the cells were fixed and stained. The expression of NP (as green fluorescence) in the Vero E6 cells was detected under a fluorescence microscope. MOCK (2% DMSO) was used as a blank control, and the position of the nucleus was indicated using DAPI staining (blue fluorescence).

### PPIX displayed its antiviral effect in the early stage of infection of influenza A virus

A time-of-addition experiment was then performed to study the detailed antiviral effect of **PPIX** on the influenza virus life cycle. In the experiment, **PPIX** was added to MDCK cells that were infected with the influenza A/PR/8/34 (H1N1) virus (100 TCID_50_) at the indicated time intervals, and the antiviral effects were then evaluated by measuring viral titers in the supernatant at 48 h postinfection. As a result, a better inhibitory effect was observed when **PPIX** was added at the early stage of infection, e.g., at the interval of (−1)-0 h and 0-2 h of treatment with the virus (Fig. 4A).

**Fig. 4.**
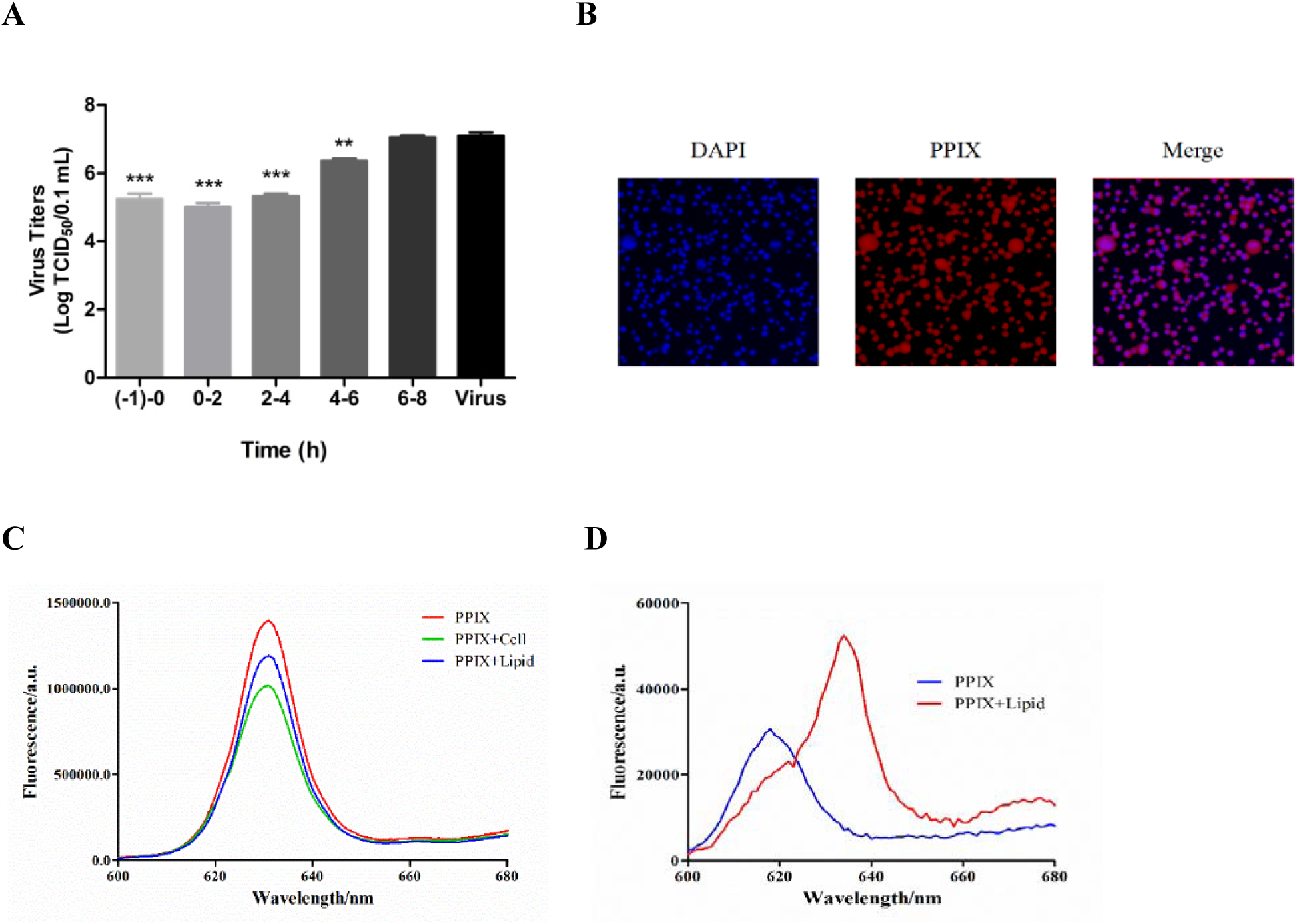
**A** Time-of-addition assay. First, 5 μg/mL **PPIX** was added to MDCK cells infected with A/PR/8/34 (H1N1) (100 TCID_50_) at the indicated time intervals. After treatment, the cells were incubated for 48 h postinfection, and the TCID_50_ values of the virus titer in the supernatant were then measured; ^*^*p*<0.05, \*\**p*<0.01, and ^***^*p*<0.001. 4**B** Fluorescence microscopic images of **PPIX** in MDCK cells. The MDCK cells in PBS were incubated with **PPIX** at a final concentration of 5 μg/mL at 37□ for 45 min, and then unbound **PPIX** was washed off twice with PBS. DAPI was used to indicate the position of the nucleus. Cells were observed under the fluorescence microscope. 4**C PPIX** at the final concentration of 5 μg/mL was added to the lipid suspension in PBS and incubated at 37□ for 45 min; then, the unbound **PPIX** was removed and the lipid was washed with PBS. The **PPIX** bound to the lipid was then extracted with 1 mL of methanol for 1 h, followed by the measurements of the fluorescence intensity at the excitation wavelength of 365 nm. **4D** Lipids were suspended in PBS at a concentration of 1 mg/mL; then, **PPIX** was added to a final concentration of 1 μg/mL. After incubation for 2 h, the fluorescence intensity in either the presence or absence of lipids was measured at the excitation wavelength of 365 nm.

This observation prompted us to further investigate the inhibitory effect of **PPIX** using four different drug administration approaches, including pretreatment of the cells, pretreatment of virus, during infection, and after infection, as reported previously (20). After drug treatment, the cells were incubated for 48 h postinfection, and then the antiviral effect was evaluated by measuring the cell survival with an MTT assay. Again, the data in Table 2 showed that the IC_50_ value of the ‘pretreatment of the virus’ approach was the most active drug administration with the IC_50_ value of 1.88±0.85 µM compared with the other approaches.

**Table 2.**
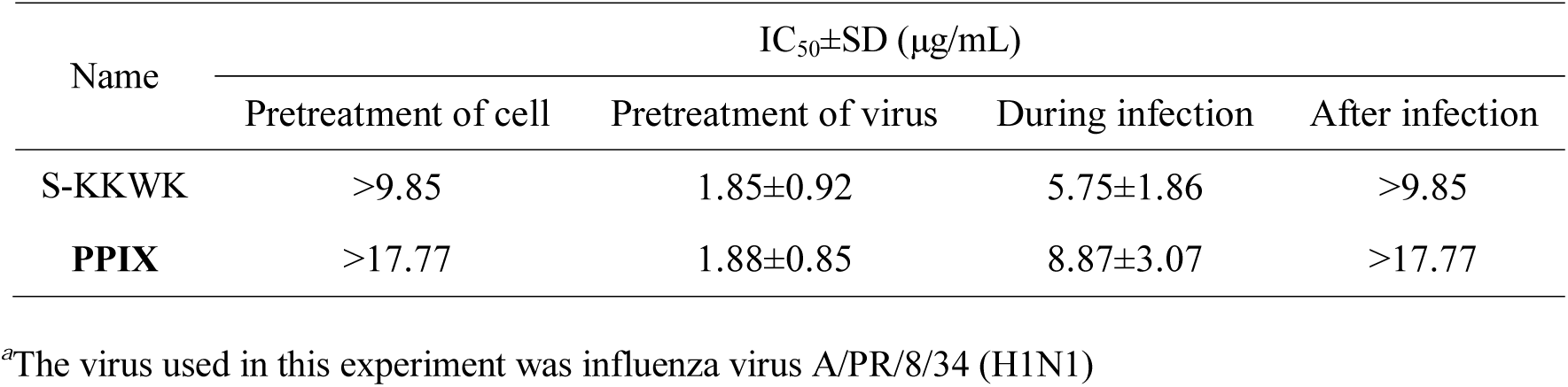
The anti-Influenza virus activity of **PPIX** using four different drug treatment approaches^*a*^

### The possible mechanism of PPIX was interaction with the lipid bilayer of enveloped virus

It is known that hemagglutinin (HA) plays a critical role in the early stage of infection by mediating the entry of IAV into host cells (23). We therefore employed two assays including the hemagglutinin inhibition assay and the hemolysis inhibition assay to study the possible interactions between HA and **PPIX** to determine whether **PPIX** can inhibit viral adsorption into target cells or inhibit the hemolytic effect on chicken erythrocytes (18). Both experiments showed negative results in the test range, indicating that hemagglutinin (HA) may not be the primary target of **PPIX** (**Fig. S1A** and **S1B**).

Due to the significant role of the viral lipid bilayer in the entry process of viruses (24), we next examined whether **PPIX** binds to the surface of the lipid bilayer. Because the lipids of the enveloped virus are derived from the host cell membrane (25), we then extracted lipids directly from MDCK cells and used them for the following study. **PPIX** at a final concentration of 5 μg/mL was added to the suspended lipid or MDCK cells in PBS and incubated for 45 min. After removal of the unbound **PPIX**, the red fluorescence of **PPIX** was detected under the fluorescence microscope (Fig. 4B), and its intensity was quantified using a fluorescence spectrophotometer at an excitation wavelength of 365 nm and a scanning range of 600 to 680 nm (Fig. 4C). **PPIX** showed apparent binding toward the lipid bilayer, as indicated in Fig. 4B and **4C.**

We further investigated the interactions between **PPIX** and lipids using fluorescence spectroscopy. The experiment was conducted by measuring the emission fluorescence from 600 to 680 nm when excited at 365 nm of 1 µg/mL of **PPIX** either in PBS or mixed with 1 mg/mL of lipids. As shown in Fig. 4D, a significant redshift from 618 to 634 nm in the maximum emission wavelength was detected when measurements were conducted in the presence of lipids.

Next, proton nuclear magnetic resonance (^1^H NMR) spectroscopic technology was used to study the interactions between **PPIX** and lipids. As indicated in **Table S1**, in the presence of lipids, the chemical shifts of a number of protons of **PPIX** moved downfield to a different extent with or without the presence of water. Moreover, the C-5, C-10, C-15 and C-20 positions of porphyrin showed more chemical shifts than the other positions (Fig. 1).

To quantitatively measure the lipid-**PPIX** interactions, isothermal titration calorimetry (ITC) technology was employed to measure the thermodynamic curves resulting from the binding of **PPIX** to lipids (26). First, 50 μg/mL **PPIX** was added to lipids at a concentration of 1 mg/mL, and the thermal changes were then measured. The thermogram in **Fig. S1C** shows that the interactions between **PPIX** and lipids were exothermic due to the negative ITC peaks, and the binding affinity K_a_ (association constant) resulting from the binding of **PPIX** to lipids was 0.01 µM^−1^ as calculated from the measured ΔH and TΔS values (**Table S2**).

### The addition of lipids reduces the anti-IAV activity of PPIX

The virus lipid bilayer as the possible target of **PPIX** prompted us to speculate whether the addition of lipids to **PPIX** would affect the anti-IAV activity of **PPIX**. The results as displayed in Fig. 5A show that the antiviral activity of **PPIX** was decreased along with the increase of lipids, further confirming the interactions between lipids and **PPIX**.

**Fig. 5.**
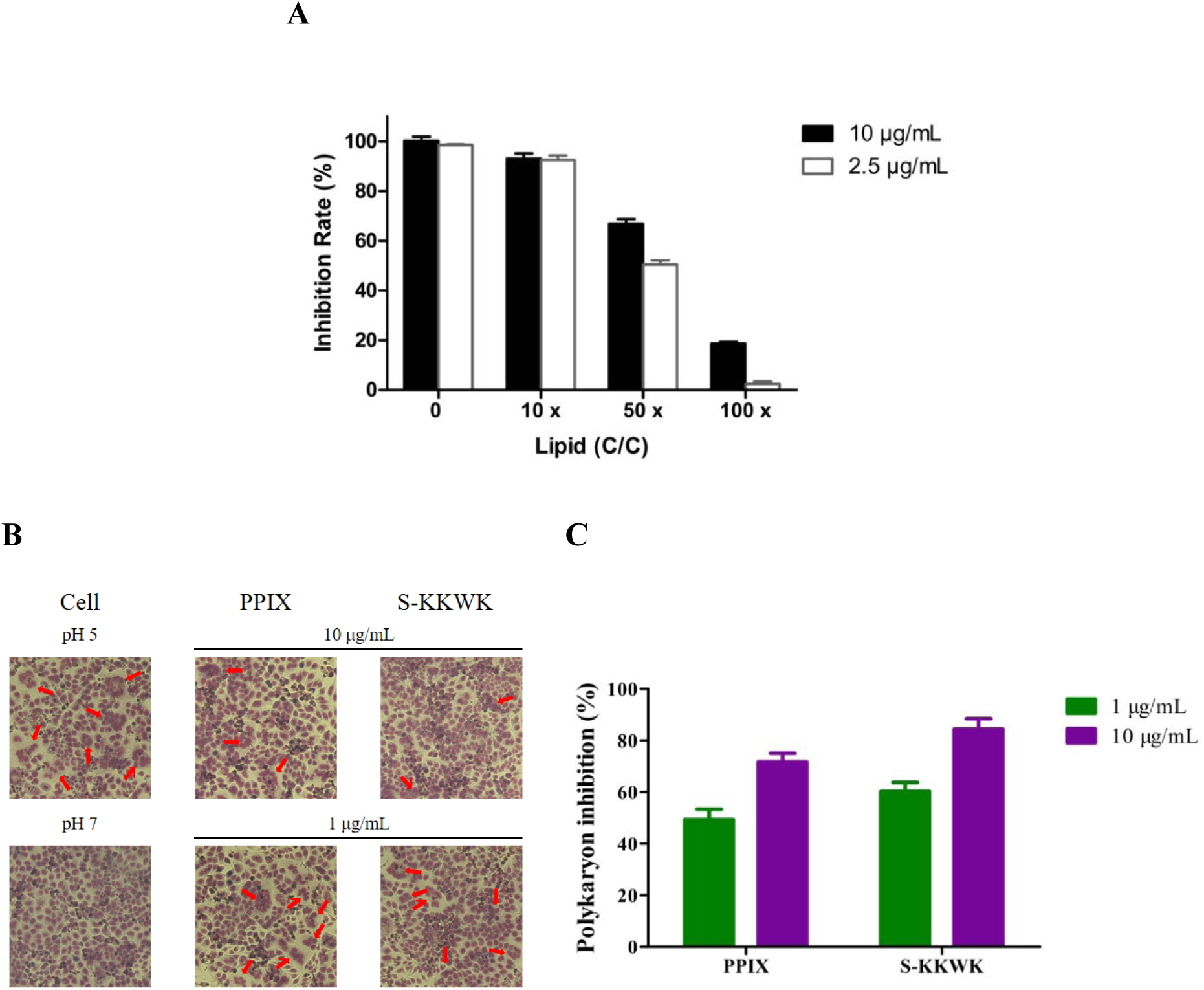
**A** The inhibition rate of **PPIX** against influenza A virus in the presence of lipids. PPIX was used at 10 and 2.5 µg/mL while the lipid concentration was 10×, 50×, and 100× that of **PPIX**. The antiviral activity was assessed using the ‘pretreatment of the virus’ approach. **5B** Polykaryon formation inhibition assay. **PPIX** or a positive control of S-KKWK was added to the MDCK cells expressing HA from influenza A/Thailand/Kan353/2004, of which the positive control was set to be the same as that reported in our previous work (17); subsequently, the culture medium was acidified to a pH of 5.0 and allowed to incubate for 15 min at 37□. After neutralization, the cells were incubated for another 3 h. Then, the cells were fixed and stained with Giemsa solution. Syncytium formation was visualized, and the number of syncytia was indicated by red arrowheads under a microscope. **5C** The number of syncytia was counted and quantified in whole fields of the plate. Each polykaryon containing at less three nuclei was counted. The syncytium formation inhibition rate was determined by comparing with the pH 5.0 control.

Due to the early inhibitory effect of **PPIX** toward the infection of influenza A virus and because the experimental data showed that **PPIX** may interact with the viral lipid bilayer to interfere with the entry of virus into host cells, we then employed a polykaryon formation inhibition assay to study the effects of **PPIX** on the formation of syncytium mediated by HA under acidic conditions (27). As shown in Fig. 5B and **5C**, the number of syncytia was significantly decreased when tested in the presence of **PPIX**, especially at a higher concentration of 10 µg/mL, where the number of syncytia was reduced by up to 80% compared with the control without drug treatment, again confirming the possible interactions between **PPIX/**HA or **PPIX/**cell membranes.

### PPIX displays antiviral activity in vivo

To explore the preliminary antiviral effect of **PPIX** *in vivo*, we next tested the anti-IAV activity of **PPIX** using mice infected with comparatively high titers of IAV. In this experiment, 4-week-old (*ca*. 19-22 g) male Kunming mice were divided into five groups with 10 mice each and intranasally challenged twice each with 2×10^5^ TCID_50_ of influenza A/PR/8/34 (H1N1) virus in a volume of 30 µL of normal saline (NS), as indicated in Fig. 6A. The drugs, either a high (10 mg/kg) or low dose (2.5 mg/kg) of **PPIX** or oseltamivir phosphate (10 mg/kg), were orally administered once a day starting from the first day after viral challenging until the fifth day. The *in vivo* antiviral effect of **PPIX** was then determined by measuring the mean number of survival days, and the survival rate was measured up to 15 days.

**Fig. 6.**
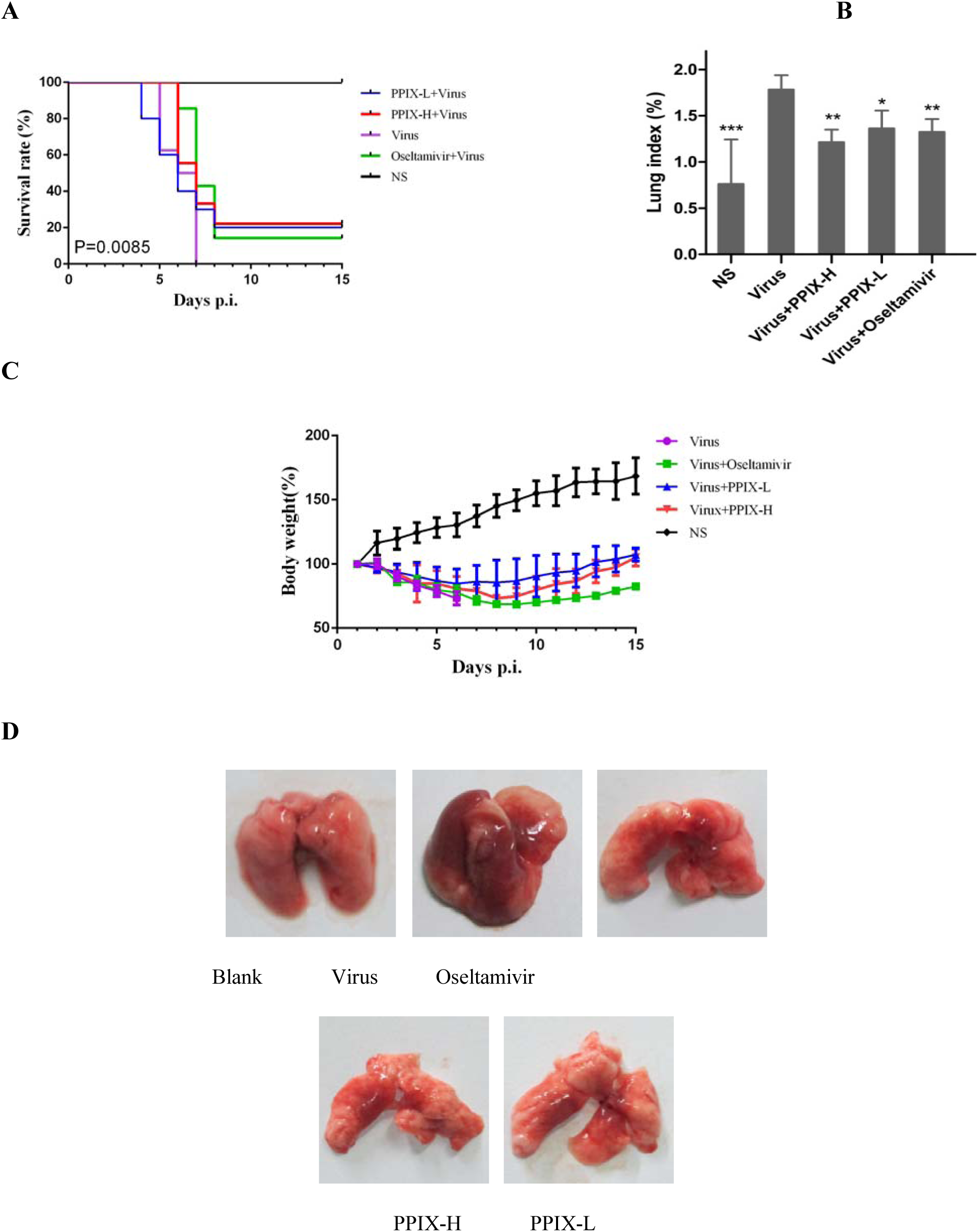
The anti-IAV activity of **PPIX** in mice. **6A** The survival rate of mice after infection with IAV and treatment with **PPIX** or oseltamivir compared with the infected mice without drug treatment. 2×10^5^ TCID_50_ of influenza virus A/PR/8/34 (H1N1) in a volume of 30 µL of normal saline (NS) was used to infect the mice. The mice were intranasally challenged with virus twice and subsequently orally administered **PPIX** (2.5 and 10 mg/kg) or oseltamivir phosphate (10 mg/kg). The drugs were administered once a day until the fifth day. The mortality rates were analyzed with GraphPad Prism 6 software by a log-rank (Mantel-Cox) test (P < 0.01). **6B** Lung index of infected mice treated with **PPIX** or oseltamivir. After the viral challenging and **PPIX** or oseltamivir treatment for 3 days, the lung index of the mice was measured. Data were analyzed with one-way ANOVA, ^*^*p*< 0.05, ^**^*p*< 0.01, and ^***^*p*< 0.001. **6C** The body weight of the infected mice was recorded. **6D** Lung images of the influenza virus-infected mice in each group after treated with different agents for three days.

The data in Fig. 6A show that the survival rate and mean survival time of the infected mice treated with high and low dosages of **PPIX** were apparently increased compared with the virus control group without drug treatment (*p*<0.01), in which all mice were dead within seven days. In addition, the lung indices measurements showed a significant increase for all virus-challenged mice compared with the control group without virus challenging (*p*<0.001). Moreover, the average lung indices for compound-treated groups were lower than for the virus control group (*p*<0.01) (Fig. 6B), which was further supported by the lung images where a dramatic difference in their lung colors was observed between compound-treated groups and virus control group (Fig. 6D).

In contrast, with the slow increase for the blank control group without virus challenging, the body weight of all virus-infected mice decreased until the eighth day, where the body weights of both the PPIX- and oseltamivir phosphate -treated groups slowly increased. Meanwhile, all mice in the virus control group died within seven days (Fig. 6C).

## Discussion

Among a number of emerging and re-emerging contagious viruses, enveloped RNA viruses account for many outbreaks (1,28), including SARS in 2003, the H1N1 epidemics in 2009, and the current COVID-19 outbreak. In addition, many viruses, such as the Lassa and Ebola viruses, that lead to severe hemorrhagic fever are highly pathogenic; thus far, there are no FDA-approved drugs available for treating these diseases (19). Therefore, the purpose of this study is to develop broad antivirals by targeting the essential components in the virus life cycle for prophylaxis and treatment of these diseases (29).

Herein, we report a porphyrin derivative of **PPIX** that showed broad anti-IAV activity against several unrelated RNA viruses, including the highly pathogenic Lassa virus (LASV) and Machupo virus (MACV) as well as SARS-CoV-2, which is responsible for the current worldwide outbreak of COVID-19. In addition, **PPIX** is also active against various subtypes of influenza A viruses (IAV), such as A/Puerto Rico/8/34 (H1N1), the A/FM/1/47 (H1N1) mouse-adapted viral strain, A/Puerto Rico/8/34 (H1N1) with the NA-H274Y mutation, and the A/Aichi/2/68 (H3N2) viral strain, with IC_50_ values ranging from 0.91±0.25 μM to 1.88±0.34 μM. These results indicate that **PPIX** may interact with a key component involved in the life cycle of these viruses.

To investigate the possible modes of action of **PPIX**, by using influenza A/Puerto Rico/8/34 (H1N1) virus as a testing strain, we employed a set of experiments involving different drug administration approaches, indirect immunofluorescence staining, qRT-PCR and plaque reduction assays (Fig. 2). The results show that the antiviral effect of **PPIX** was mainly observable in the early stage of infection. Based on these results, we further identified that the activity may have resulted from the interactions of **PPIX** with the lipid bilayer of the virus envelope, thereby inhibiting the entry of virus; this was deduced from fluorescence spectroscopy (Fig. 3B, Fig. 4A and **Fig. S1A**), NMR experiments (**Table S1**), and semiquantitative ITC analyses (**Table S2** and **Fig. S1C**). This conclusion is consistent with the results of our previous study, showing that the antiviral activity of Pyropheophorbide a (**PPa**) was due to the interaction with the viral membrane or biophysical interference with the virus-cell membrane fusion process, which resulted in blocking the entry of an enveloped virus into cells (17).

Nevertheless, considering the pivotal role of hemagglutinin (HA) of influenza A virus in mediating the entry of the virus, the results of the fusion inhibition assay (Fig. 4C and **4D**) cannot exclude the possibility that the antiviral activity may have resulted from the interaction of **PPIX** with the surface glycoprotein HA, blocking the entry of the virus.

However, (1) both the negative results of the hemagglutinin inhibition assay and hemolysis inhibition assay (Fig. 2D and **2E**) showed that **PPIX** was unable to either block the absorption of virus onto host cells or inhibit the fusion of virus with the host cell membrane (18,22); (2) **PPIX** showed broad antiviral activities against a number of unrelated enveloped viruses belonging to different families; and (3) **PPIX** showed interactions with the lipid bilayer. Therefore, we reasonably deduced that the major antiviral properties were not mediated by a specific receptor of the viral particles. Instead, the observed inhibitory effect in the fusion inhibition assay (Fig. 4C) may be attributed to the interaction of **PPIX** with the lipid bilayer.

From the structural point of view (Fig. 1), **PPIX** possesses a rigid amphipathic skeleton with two polar carboxy groups and a nonpolar moiety of aromatic rings. These features enable **PPIX** to intercalate with hydrophobic lipids of enveloped virions or cell membrane (30,31), thereby inhibiting the formation of the negative curvature required for fusion (32) or inhibiting virion-cell lipid mixing (30); as a result, these features inhibit infectivity of a broad variety of unrelated viruses^31^. When **PPIX** was modified with a large nonpolar group, as indicated in Fig. 1, the structural integrity as well as the amphipathicity of **PPIX** was compromised, correspondingly diminishing the antiviral effectivity.

With respect to the selective virion-to-cell activity of **PPIX** over that of cellular fusion, based on the model proposed by Colpitts *et al*, it could be deduced that **PPIX** functions by increasing the hemifusion stalk energy barrier when intercalated in lipids (30,31). Compared with virions that are metabolically inert, cells are able to use metabolic energy to reshape lipids and bring membranes together. These different properties between virions and cells may lead to the observed selectivity of **PPIX** when interacting with virions and host cells (30,32–34). Therefore, it can be deduced that the fusion inhibitory activity of **PPIX** resulted from the biophysical interactions rather than the chemical interactions with lipids (32). However, the proposed mechanism remains to be investigated.

In this study, we report the antiviral properties of **PPIX**. Structurally, it possesses a porphyrin core with two aliphatic carboxy groups; therefore, **PPIX** is a new rigid amphipathic fusion inhibitor (RAFIs) (33). It showed broad-spectrum antiviral activities toward a panel of unrelated enveloped viruses, including highly pathogenic Lassa virus (LASV) and Machupo virus (MACV) as well as SARS-CoV-2. The mechanism study indicated that **PPIX** biophysically interacted with the lipid bilayer of the enveloped virus, inhibiting the membrane-associated functions required for virion-to-cell fusion. Because membrane fusion is an essential step for most enveloped viruses, **PPIX** showed a broad spectrum of antiviral activity toward many unrelated enveloped viruses, Nevertheless, detailed information on how the interaction with the lipid bilayer inhibits the fusion process is still unknown. Even so, **PPIX** exerts broad antiviral activity and can still be used as a lead compound for further rational modification to obtain more potent candidates for the treatment of infections with enveloped viruses.

## Materials and Methods

### Anti-influenza A virus assay

The protocol was adopted as previously reported^19^. MDCK cells were seeded in 96-well plates of 100 μL/well (2 × 10^4^ cells) and cultured in a 5% CO_2_ incubator at 37℃ for 24 h. Influenza A/Puerto Rico/8/34 (H1N1), A/FM/1/47 (H1N1), A/Puerto Rico/8/34 (H1N1) with the NA-H274Y mutation, and A/Aichi/2/68 (H3N2) viruses at 100 TCID_50_ were each mixed with twofold-diluted PPIX solutions and incubated at 37°C for 30 min. After washing twice with PBS (Sigma, USA), the virus-compound mixtures were added to these cells and incubated for another 1 h. After that, DMEM (Sigma, USA) supplemented with 1 μg/mL of TPCK-trypsin (Sigma, USA) was added to the MDCK cells. At 48 h postinfection, cell viability was measured by the MTT method. S-KKWK was used as a positive control (20), and the experiment was repeated at least three times independently.

### Anti-Lassa or Machupo pseudotyped viruses assay

Vero-E6 cells were seeded in 96-well plates with 1.5×10^4^ cells/well overnight. Lassa or Machupo pseudotyped viruses were incubated with gradient-diluted compounds at 37℃ for 30 min, then the mixture was added to cells and incubated at 37℃ for 1 h. After that, the supernatants were completely removed, cells were washed with PBS and cultured for 24 h before being subjected to luciferase activity detection (19).

### Anti-SARS-CoV-2 assay

Vero-E6 cells were seeded in 48-well plates with 5×10^4^ cells/well overnight. For ‘pretreatment of virus’ mode, SARS-CoV-2 were processed as the same way as Lassa or Machupo pseudotyped viruses. For ‘full-time treatment’ mode, gradient-diluted compounds were firstly, incubated with cells for 1 h, then SARS-CoV-2 were added and incubated with cells for another 1 h. Then the supernatants were completely removed, cells were washed with PBS and supplemented with medium containing gradiently-diluted compounds. After 24 h, cell supernatants were collected and subjected to viral RNA isolation, then viral genome copies were detected by qRT-PCR with primers targeting S gene. At the same time, cells were fixed with 4% formaldehyde and viruses were detected by indirect immunofluorescence with polyantibodies against NP (22).

### Polykaryon formation inhibition assay

Following a procedure previously reported by Lai et al. (27), Briefly, MDCK cells (2 ×10^5^ cells per well; 12-well plate) incubated for 16 h. According to the manufacturer’s instructions transfected with plasmid-encoded HA (from influenza A/Thai/Kan353/ 2004 virus) (2 μg DNA/well) using PEI. After 8-10 h of culture, fresh DMEM containing 10% fetal bovine serum was used instead of transfection medium. At 48 h after transfection, the cells were washed twice with PBS and treated with TPCK-trypsin (5 μg/mL) for 15 min at 37℃. Then, the cells were rinsed with PBS solution twice and pretreated with 0.5 mL of **PPIX** or 1% methanol for 15 min at 37℃ followed by an incubation with 0.5 mL of pH 5.0 PBS or 1% methanol at 37℃ for another 15 min. After the reaction, the cells were washed twice with PBS, added 1 mL of DMEM containing 10% FBS incubated at 37℃ to form polykaryon. 3 h later, the cells were fixed with methanol and incubated with Giemsa stain (Sigma, USA), and visualized under a microscope. Syncytium formation was quantified by counting the number of polynuclei (containing 5 or more nuclei) under a microscope, and the percentage of syncytium formation was determined relative to methanol. S-KKWK was used as a positive control, and the experiment was repeated at least three times independently.

### Anti-influenza virus test in vivo

#### Animals

Four-week-old male KM mice without a specific pathogen (SPF) with an average body weight of 19-24 g were purchased from the Laboratory Animal Center of Southern Medical University (SMU) (Guangzhou, China). All mice were fed standard laboratory chow and provided *ad libitum* access to water during the animal experiments. The experiments were monitored in accordance with the Standard Operating Procedures of the facility and the Animal Welfare Act.

The mice were randomly divided into five groups, including the uninfected and water-treated group (blank), the infected and water-treated group (viruses), the group that was infected and treated with 10 mg/mL oseltamivir phosphate, and the groups that were infected and treated with **PPIX** at concentrations of 10 mg/kg (**PPIX**-H) and 2.5 mg/kg (**PPIX**-L). Oseltamivir phosphate was dissolved in water, whereas **PPIX** was initially dissolved in polyethylene glycol 400 and then diluted fourfold with water. Infected mice were anesthetized with ethyl ether and then intranasally challenged twice with influenza A/PR/8/34 (H1N1) virus at the titer of 2×10^5^TCID_50_ in a volume of 30 μL. The mice then took a different drug orally once a day for five days. The mice were given food and water *ad libitum* on the sixth day. Body weight, mortality, and the general behaviors of the mice were recorded for fifteen consecutive days.

#### Lung index

Using the same protocol as the *in vivo* anti-influenza virus test described above, on the fourth day after infection drug administration to the stomach was stopped in each group of three mice, and the animals were sacrificed. Their lungs were harvested, washed with normal saline, dried with gauze, and weighed. Meanwhile, the lung images were obtained (Fig. 6D). The lung index was calculated using the following equation:

Lung index = lung weight/body weight × 100%

#### Statistical analysis

The half-cytotoxic concentration (CC_50_) and half-inhibitory concentration (IC_50_) values of the PPIX were calculated with GraphPad Prism 5 software (San Diego, CA). Each data point, expressed as the means ± standard deviation (SD), was repeated at least three to five times. Fluorescent images were greyscale quantized using ImageJ. The mortality rates were analyzed with GraphPad Prism 6 software by a log-rank (Mantel-Cox) test. Lung index were determined by one-way ANOVA using SPSS 22.0 software. Statistical significance was defined as ^*^*p*< 0.05, \*\**p*< 0.01, ^***^*p*< 0.001.

## Acknowledgments

The authors wish to thank Dr. Guang Yang for his measurements of ITC data. This work was funded by the National Natural Science Foundation of China (81773556), China, Science and Technology Department of Guangdong Province of China (2015A020211010), and Southern Medical University (B1040903).

## Author contributions

S. S. L, X. Y. P, D. W. C, Y. W, W. J. S, X. M. J, Y. S. performed the experiments and part of data analyses. S. F. assisted in the animal experiment and prepared the figures. J. H. conceived and designed the research and wrote the manuscript. All authors contributed to review of this manuscript.

## References and Notes

1. Howard CR, Fletcher NF. Emerging virus diseases: can we ever expect the unexpected? Emerg. Microbes. Infect. 2012;1, e46.

2. Smith AE, Helenius A. How viruses enter animal cells. Science. 2004;304, 237–242.

3. Marsh M, Helenius A. Virus entry: Open sesame. Cell. 2006;124, 729–740.

4. Yamauchi Y, Helenius A. Virus entry at a glance. J. Cell. Sci. 2013;126, 1289–1295.

5. Mercer J, Schelhaas M, Helenius A. Virus Entry by Endocytosis. Annu. Rev. Biochem. 2010;79, 803–833.

6. Edinger TO, Pohl MO, Stertz S. Entry of influenza A virus: host factors and antiviral targets. J. Gen. Virol. 2014;95, 263–277.

7. Martens S, Mcmahon HT. Mechanisms of membrane fusion: disparate players and common principles. Nat. Rev. Mol. Cell. Biol. 2008;9, 543–556.

8. Badani H, Garry RF, Wimley WC. Peptide entry inhibitors of enveloped viruses: the importance of interfacial hydrophobicity. Biochim. Biophys. Acta. 2014;1838, 2180–2197.

9. Bobardt MD, Cheng G, de Witte L, Selvarajah S, Chatterji U, Sanders-Beer BE, Geijtenbeek TB, Chisari FV, Gallay PA. Hepatitis C virus NS5A anchor peptide disrupts human immunodeficiency virus. Proc. Natl. Acad. Sci. U S A. 2008;105, 5525–5530.

10. Cheng G, Montero A, Gastaminza P, Whitten-Bauer C, Wieland SF, Isogawa M, Fredericksen B, Selvarajah S, Gallay PA, Ghadiri MR, Chisari FV. A virocidal amphipathic α-helical peptide that inhibits hepatitis C virus infection *in vitro*. Natl. Acad. Sci. U S A. 2008;105, 3088–3093.

11. Mody TD. Pharmaceutical development and medical applications of porphyrin-type macrocycles. J. Porphyr. Phthalocya. 2000;4, 362–367.

12. Ballut S, Naud-Martin D, Loock B, Maillard P. A Strategy for the targeting of photosensitizers. Synthesis, characterization, and photobiological property of porphyrins bearing glycodendrimeric moieties. J. Org. Chem. 2011;76, 2010–2028.

13. Chen S, Fetzer JC, Meyerhoff ME. Pharmaceutical development and medical applications of porphyrin-type macrocycles. Fresenius. J. Anal. Chem. 2001;369, 385–392.

14. Armstrong NR. Phthalocyanines and porphyrins as materials. J. Porphyr. Phthalocya. 2000;4, 414–417.

15. Choi SR, Britigan BE, Narayanasamy P. Dual inhibition of *Klebsiella pneumonia* and *Pseudomonas aeruginosa* iron metabolism using gallium porphyrin and gallium nitrate. ACS. Infect. Dis. 2019;5, 1559–1569.

16. Wang XB, Wang Y, Wang P, Cheng XX, Liu QH. Sonodynamically induced anti-tumor effect with protoporphyrin IX on hepatoma-22 solid tumor. Ultrasonics. 2010;51, 539–546.

17. Chen D, Lu S, Yang G, Pan X, Fan S, Xie X, Chen Q, Li F, Li Z, Wu S, He J. The seafood *Musculus senhousei* shows anti-influenza A virus activity by targeting virion envelope lipids. Biochem Pharmacol. 2020. doi: 10.1016/j.bcp.2020.113982.

18. Lin DG, Li FF, Wu QY, Xie XK, Wu WJ, Wu J, Chen Q, Liu SW, He J. A ‘building block’ approach to the new influenza A virus entry inhibitors with reduced cellular toxicities. Sci. Rep.2016;6, 22790.

19. Wang PL, Liu Y, Zhang GS, Wang SB, Guo J, Cao JY, Jia XY, Zhang LK, Xiao GF, Wang W. Screening and identification of Lassa virus entry inhibitors from an FDA-approved drug library. J. Virol. 2018;92, e00954–18.

20. Lin DG, Luo YZ, Yang G, Li FF, Xie XK, Chen DW, He LF, Wang JY, Ye CF, Lu SS, Lv L, Liu SW, He J. Potent influenza A virus entry inhibitors targeting a conserved region of hemagglutinin. Biochem. Pharmacol. 2017;144, 35–51.

21. Zu M, Yang F, Zhou WL, Liu AL, Du GH, Zheng LS. *In vitro* anti-influenza virus and anti-inflammatory activities of theaflavin derivatives. Antiviral. Res. 2012;94, 217–224.

22. Wang ML, Cao RY, Zhang LK, Yang XL, Liu J, Xu MY, Shi ZL, Hu ZH, Zhong W, Xiao GF. Remdesivir and chloroquine effectively inhibit the recently emerged novel coronavirus (2019-nCoV) *in vitro*, Cell. Res. 2020;30, 269–271.

23. Gamblin SJ, Haire LF, Russell RJ, Stevens DJ, Xiao B, Ha Y, Vasisht N, Steinhauer DA, Daniels RS, Elliot A, Wiley DC, Skehel JJ. The structure and receptor binding properties of the 1918 influenza hemagglutinin. Science. 2004;303, 1838–1842.

24. Wu WJ, Lin DG, Shen XT, Li FF, Fang YX, Li KQ, Xun TR, Yang G, Yang J, Liu SW, He J. New influenza A virus entry inhibitors derived from the viral fusion peptides. Plos. One. 2015;10, e138426.

25. Kuhn RJ, Strauss JH. Enveloped viruses. Adv. Protein. Chem. 2003;64, 363–377.

26. Myszka DG. Kinetic, equilibrium, and thermodynamic analysis of macromolecular interactions with BIACORE. Method. Enzymol. 2000;323, 325–340.

27. Lai KK, Cheung NN, Yang F, Dai J, Liu L, Chen ZW, Sze KH, Chen HL, Yuen KY, Kao RYT. Identification of novel fusion inhibitors of influenza A virus by chemical genetics. J. Virol. 2015;90, 2690–2701.

28. Yee HY, Luo DH. RIG-I-Like receptors as novel targets for pan-antivirals and vaccine adjuvants against emerging and re-emerging viral infections. Front. Immunol. 2018;9, 1379.

29. Halfon P, Sarrazin C. Future treatment of chronic hepatitis C with direct acting antivirals: is resistance important? Liver. Int. 2012;1:79–87.

30. Colpitts CC, Ustinov AV, Epand RF, Epand RM, Korshun VA, Schang LM. 5-(Perylen-3-yl) ethynyl-arabino-uridine (aUY11), an arabino-based rigid amphipathic fusion inhibitor, targets virion envelope lipids to inhibit fusion of influenza virus, hepatitis C virus, and other enveloped viruses. J. Virol. 2013;87, 3640–3654.

31. St Vincent MR, Colpitts CC, Ustinov AV, Muqadas M, Joyce MA, Barsby NL, Epand RF, Epand RM, Khramyshev SA, Valueva OA, Korshun VA, Tyrrell DL, Schang LM. Rigid amphipathic fusion inhibitors, small molecule antiviral compounds against enveloped viruses. Proc. Natl. Acad. Sci. U S A. 2010;107, 17339–17344

32. Speerstra S, Chistov AA, Proskurin GV, Aralov AV, Ulashchik EA, Streshnev PP, Shmanai VV, Korshun VA, Schang LM. Antivirals acting on viral envelopes via biophysical mechanisms of action. Antiviral. Res. 2018;149, 164–173.

33. Vigant F, Hollmann A, Lee J, Santos NC, Jung ME, Lee B. The rigid amphipathic fusion inhibitor dUY11 acts through photosensitization of viruses. J. Virol. 2014;88, 1849–1853.

34. Vigant F, Santos NC, Lee B. Broad-spectrum antivirals against viral fusion. Nat. Rev. Microbiol. 2015;13, 426–437.

