## Supplementary Materials for "Broad-spectrum antivirals of protoporphyrins inhibit the entry of highly pathogenic emerging viruses"

**Table of contents**

Materials and Methods	3

Figure legends	11

Fig. S1 Hemagglutination inhibition assay, hemolysis inhibition assay and ITC assay of **PPIX**	12

Table legends	13

Table S1 Changes of ^1^H NMR chemical shift after binding between **PPIX** and MDCK lipid	14

Table S2 Thermodynamic parameters of lipid interaction between **PPIX** and MDCK cells	15

**Materials and Methods**

***Chemicals and analytical instruments***

NMR spectra were acquired with a Bruker DRX-400 spectrometer (400 MHz for ^1^H NMR, 100 MHz for ^13^C NMR, respectively). Chemical shifts were referenced to the corresponding residual solvent peak as internal standard (7.26/77.23 ppm in CDCl_3_). Low-resolution ESI-MS were recorded on a Waters 3100 single-quadrupole mass spectrometer. All ragents were obtained from commercial sources and used without further treatment. Silica gel (300-400 mesh, Qingdao Marine Chemical Factory, China), Sephadex LH-20 (Amersham Pharmacia Biotech) were used for the column chromatography. Positive control of oseltamivir phosphate was purchased from Yichang Dongyangguang Pharmaceutical Co., Ltd.

***General procedure for synthesis and purification of PPIX-1~5***

**PPIX** was dissolved in DMF and the solution was allowed to cool at 0℃ with ice bath. Then the corresponding alcohols dissolved in DMF were added into the reaction mixture, followed with the reagents including HBTU, DMAP, and DCC. HBTU and DMAP were dissolved in DMF while DCC was in dichloromethane. DIEA was subsequently added to the reaction mixture, which was then allowed to stir at room temperature until the end of the reaction, where **PPIX** was disappeared and a new product was observed on TLC under UV detector at 365nm.

The product solution was dissolved with equivalent deionized water and extracted with ethyl acetate (3 × 25 mL). The organic phase was combined together and the solvent was evaporated under reduced pressure. The residue was purified with column chromatography on Sephadex LH-20 (with MeOH as eluent) and silica gel (eluent system: dichloromethane-methanol) to give corresponding products (**Fig.1**). **PPIX-1**: prepared from 1-decanol (28.4 mg). Yielded: 8.42 mg. **PPIX-2**: prepared from 1-tetradecanol (9.6 mg). Yielded: 3.47 mg. **PPIX-3**: prepared from 1-octadecanol (15.1 mg). Yielded: 5.5 mg. **PPIX-4**: prepared from benzyl alcohol (4.61 mg). Yielded: 3.4 mg. **PPIX-5**: prepared from 4-hydroxymethylbiphenyl (7.89 mg). Yielded: 7.2 mg.

***Cells and viruses***

African green monkey kidney Vero E6 cells (ATCC1586) and Madin Darby Canine Kidney (MDCK) cells were obtained from the American Type Culture Collection (ATCC). MDCK cells were maintained in Dulbecco’s Modified Eagle Medium (DEME, Sigma, USA) containing glutamine, supplemented with 10% fetal bovine serum (FBS, Sigma, USA) and 1% penicillin/streptomycin, and Vero E6 cells were cultured in minimum Eagle’s medium (MEM, Sigma, USA) supplemented with 10% FBS. Both cells were grown at 37℃ in a humidified atmosphere of 5% CO_2_.

Influenza A subtypes including A/Puerto Rico/8/34 (H1N1), A/FM/1/47 (H1N1) mouse-adapted viral strain, A/Puerto Rico/8/34 (H1N1) with the NA-H274Y mutation, and A/Aichi/2/68 (H3N2) viral strains were propagated in 9-day-old embryonated hen eggs at 37℃, and the allantoic fluid containing above viruses were stored at -80℃. The viral titer was measured in 50% tissue culture infectious dose (TCID_50_) test.

Vesicular stomatitis virus (VSV) backboned-Lassa and Machupo pseudotyped viruses were packaged as previously described (1). In brief, plasmid vectored with glycoprotein gene of Lassa (GenBank: NC_004296.1) or Machupo (GenBank: NC_005078.1) was transfected into HEK293T cells. 24 h later, cells were infected with VSVΔG-GFP/VSV G (packaged by our laboratory members) for 1 h, then the cell supernatant was completely removed, and culture supernatants containing pseudovirus (VSVΔG-GFP/Lassa G or VSVΔG-Rluc/MACV G) were collected after 36 h. These pseudoviruses were restricted to a single round of replication, thus enabling them to be handled in a BSL-2 lab.

SARS-CoV-2 was preserved at Wuhan institute of Virology and handled in BSL-3 lab. It was propagated in Vero-E6 cells and calculated copy numbers by qRT-PCR before use (2).

***Antiviral assay with different drug administration protocols***

As with a previous study (3), four different modes of administration were used to explore the possible mechanism of **PPIX**. (1) Pretreatment of cells: PPIX was added to cell monolayers for 30 min before influenza virus A/PR/8/34 (H1N1) (100 TCID_50_) adsorption at 37°C. (2) Pretreatment of virus: **PPIX** was preincubated with influenza virus A/PR/8/34 (H1N1) for 30 min at 37°C before being added to the MDCK cells. (3) During infection: A mixture of **PPIX** and influenza virus A/PR/8/34 (H1N1) was added to the cells simultaneously. (4) After infection: **PPIX** was added after the influenza virus A/PR/8/34 (H1N1) (100 TCID_50_) adsorbed onto the cells. The cells were washed with PBS and cultured with serum-free DMEM containing 1 μg/mL TPCK-trypsin for 48 h. The cell survival rate was determined by MTT assay after 48 h. The cytopathological effect (CPE) of virus-infected cells was observed under a microscope.

***Plaque reduction assay***

MDCK cells were cultured in 6-well plates of 2 mL/well (4×10^5^ cells) for 24 h until confluent. Antiviral effects were evaluated by different drug administration protocols: ‘During infection’ and ‘Pretreatment of virus’, as mentioned above (3). After virus absorption, inoculums were removed and cells were washed twice with PBS, then 4 mL of serum-free MEM (2×) containing 1μg/mL TPCK-trypsin (Sigma, USA) (1.6% AGAR for 72 h, as mentioned above) was added into cells. After infection for 72 h, the cell monolayer was fixed with 4% paraformaldehyde for 20 min. Then agarose overlay was removed and cell monolayer was stained with 0.5% (w/v) crystal violet dye solution for 1 h at 37℃. Through counting number of plaques, determine the PPIX effect on virus plaque formation.

***Quantitative real-time PCR assay***

4×10^5^ MDCK cells per well was seeded into 6-well plates were subjected to two different modes of administration including ‘during infection’ and ‘pretreatment virus’. 24 h after infection, the total RNA was extracted using Trizol reagent (Sigma, St. Louis, MO, USA), and the primers listed below were used to reverse transcribed the RNA into cDNA. Two-step PCR amplification standard procedure of ABI7500 system (Applied Biosystems, Foster, CA, USA) was used for real-time quantitative PCR. Glyceraldehyde 3-phosphate dehydrogenase (GAPDH, Sigma, USA) was used as an internal reference. After that, the relative content of the HA gene was compared by 2^−ΔΔCT^ method. Each sample was tested at least three times independently. The primer sequences of target genes are as follows: 5’-TTCCCAAGATCCATCCGGCAA-3’ (HA-Forward), 5’-CCTGCTCGAAGACAGCCACAACG-3’(HA-Reverse), 5’-AGG GCAATGCCAGCCCCAGCG-3’(GAPDH-Forward), 5’-AGGCGTCGGAGGGCCC CCTC-3’ (GAPDH-Reverse).

***Immunofluorescence staining and microscopy***

MDCK cells were cultured in 48-well plates and then infected with influenza virus A/PR/8/34 (H1N1) at 100 TCID_50_ using ‘Pretreatment of virus’ and ‘During infection’ protocols as described above. 24 h later, after removal of the supernatant, the cells were washed twice with PBS and then were fixed with 4% paraformaldehyde for 20 min at room temperature. Then the cells were washed three times with PBS, blocked with 4% bovine serum albumin (BSA) for 4 h at 37℃, and then incubated with nucleoprotein (NP) antibody (1:250 diluted in 4% BSA; Santa Cruz, Texas, USA) overnight at 4℃, followed by an incubation with a secondary antibody conjugated with fluorescein isothiocyanate (FITC) (1:250 diluted in 4% BSA) for another 4 h at room temperature. Then, cells were counterstained with 4,6-diamino-2-phenylindole (DAPI, Sigma, USA) for 10 min after washed three times with PBS, and the plate was monitored under a fluorescence microscope (ZEISS Axio Observer, Germany).

***Time-of-Addition assay***

MDCK cells were seeded in 6-well plates at 4 × 10^5^ cells for each well, and cultured at 5% CO_2_ for 24 h at 37℃ until the cells were monolayered. After that, the medium was discarded, washed twice with PBS and then incubated with 1 mL of influenza virus solution (100 TCID_50_) for 1 h, Then 5 μg/mL of **PPIX** was added at different time intervals (0-2 h, 2-4 h, 4-6 h, 6-8 h , respectively) during the single round replication of the virus. After 24 h, the cells were frozen and thawed repeatedly three times until the cells were completely disrupted, the supernatant was collected, the cell debris was removed by centrifugation, and the virus titer was measured by the TCID_50_ method.

***Fluorescence spectroscopy analyses***

The collected MDCK cells were washed twice with PBS and then suspended in 10 mL PBS. Before shaking for 90 min, 20 mL of chloroform-methanol (1:2, v/v) was added. Then, 20 mL of chloroform: water (1:1, v/v) was added, and the mixture was shaken well for another 30 min. The chloroform phase was separated by a separation funnel. The solvent was removed with a rotary evaporator, and the lipid was obtained.

A certain amount of PBS was added to 100 μg/mL of lipid, and the lipid suspension was obtained by ultrasound. Then, 1 mL of aliquot was transferred to several Eppendorf tubes to measure the binding of **PPIX** to a lipid bilayer. **PPIX** with a final concentration of 5 μg/mL was added to the lipid suspension. After the mixture was thoroughly mixed, the mixture was incubated at 37℃ for 45 min; the supernatant was removed by centrifugation and washed twice with PBS. Then, 1 mL of methanol was added to the lipid and shaken for 20 min to dissolve the **PPIX** absorbed on the lipids. The fluorescence intensity was then measured with a fluorescence spectrometer (FluoroMax-4, HORIBA, France) at an excitation wavelength of 365 nm and a scanning range of 500~800 nm.

The interaction between **PPIX** and lipids was further investigated by measuring the fluorescence intensity. The extracted lipids were mixed with PBS at a concentration of 1 mg/mL, and **PPIX** was added to achieve a final concentration of 1 μg/mL. The samples were incubated in the dark for 2 h, detected by fluorescence emission spectra, and compared with the sample lacking lipids.

***Isothermal titration calorimetry (ITC) assay***

Lipids extracted from MDCK cells were dissolved in 5% DMSO at a concentration of 1mg/mL and degassed for 10 min. The caloric value of PPIX (75 g/mL) interacting with lipids was recorded by isothermal titration calorimetry (MicroCal PEAQ -ITC, Malvern, UK). The ITC reaction conditions were set as follows: Temperature: 25℃; Number of injections: 19; Volume: 2 μL; Spacing: 100 s; Reference power: 10 μcal/s; Stirring speed: 750 rpm. The MicroCal peak-ITC analysis software was used to analyze the experimental data and provide a thermal parameter calculation model. The average MW of lipids was set to be 3,000 (4).

**References**

1. Pan XY, Wu Y, Wang W, Zhang LK, Xiao GF. Development of horse neutralizing immunoglobulin and immunoglobulin fragments against Junín virus. *Antiviral. Res*. 2020;174, 104666.

2. Wang ML,  Cao RY,  Zhang LK,  Yang XL,  Liu J,  Xu MY,  Shi ZL,  Hu ZH,  Zhong W,  Xiao GF. Remdesivir and chloroquine effectively inhibit the recently emerged novel coronavirus (2019-nCoV) *in vitro*, *Cell. Res*. 2020;**30**, 269–271.

3. Lin DG, Luo YZ, Yang G, Li FF, Xie XK, Chen DW, He LF, Wang JY, Ye CF, Lu SS, Lv L, Liu SW, He J. Potent influenza A virus entry inhibitors targeting a conserved region of hemagglutinin. *Biochem. Pharmacol*. 2017;**144**, 35–51.

4. Yang G, Wang JY, Lu SS, Chen Z, Fan S, Chen DW, Xue HX, Shi WY, He J. Short lipopeptides specifically inhibit the growth of Propionibacterium acnes with a dual antibacterial and anti-inflammatory action. *Br. J. Pharmacol*. 2019;**176**, 2321–2335.

**Figure legends**

**Fig. S1** Hemagglutination inhibition assay, hemolysis inhibition assay and ITC assay of **PPIX**. S**1 A** Hemagglutination inhibition (HI) assay to assess the inhibition of **PPIX** on the absorption of the virus to target cells, as described previously *(21)*. **S1 B** Hemolysis inhibition assay. A freshly prepared chicken erythrocytes suspension (2%, v/v) in PBS was mixed with **PPIX** (15 μg/mL) and an equal volume of influenza virus A/PR/8/34 (H1N1) allantoic fluid (10^6^ TCID_50_/0.1 mL). The mixture was then acidified to a pH of 4.6 to 6.0 (sodium acetate, 0.5 M) and then allowed to incubate at 37℃ for 30 min. Subsequently, the mixture was centrifuged, and the supernatants containing released hemoglobin were measured at OD_535_ using a Multiskan FC microplate reader. S-KKWK at 15 μg/mL was used as a positive control *(21)*. **S1 C** The thermogram of the binding of 50 μg/mL **PPIX** to 1 mg/mL of lipids was obtained by an ITC assay. The thermal changes due to the interactions between **PPa** and cellular lipids were detected.

**A	B**

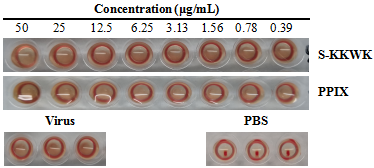

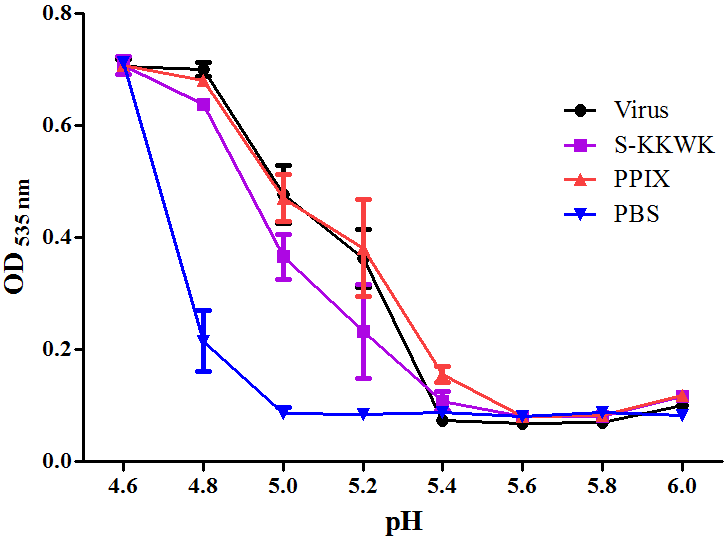
 **C**

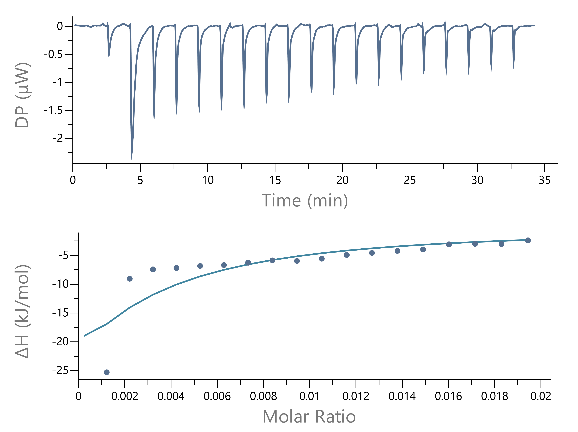

**Fig. S1**

**Table legends**

**Table S1** Changes of ^1^H NMR chemical shift after binding between **PPIX** and MDCK lipid

**Table S2** Thermodynamic parameters of lipid interaction between **PPIX** and MDCK cells

**Table S1** Changes of ^1^H NMR chemical shift after binding between **PPIX** and MDCK lipid

| Position | *δ* _H_ ^a^ | | Δ*^b^* | *δ* _H_ | | Δ*^c^* | *δ* _H_ | | Δ*^d^* |
| --- | --- | --- | --- | --- | --- | --- | --- | --- | --- |
|  | **PPIX** | **PPIX**+Lipid |  | **PPIX**  (10% H_2_O) | **PPIX**+Lipid  (10% H_2_O) |  | **PPIX**  (16.67% H_2_O) | **PPIX**+Lipid  (16.67% H_2_O) |  |
| 5/10/15/20 | 10.333 | 10.372 | 0.039 | 10.210 | 10.259 | 0.049 | 10.010 | 10.121 | 0.111 |
| 5/10/15/20 | 10.267 | 10.296 | 0.029 | 10.165 | 10.209 | 0.044 | 9.977 | 10.088 | 0.111 |
| 3’/8’ | 8.517 | 8.535 | 0.018 | 8.447 | 8.469 | 0.022 | 8.353 | 8.400 | 0.047 |
| 3’’/ 8’’ | 6.468 | 6.480 | 0.012 | 6.443 | 6.459 | 0.016 | 6.404 | 6.435 | 0.031 |
| 3’’/ 8’’ | 6.233 | 6.239 | 0.006 | 6.241 | 6.253 | 0.012 | 6.231 | 6.256 | 0.025 |
| 13’, 17’ | 4.351 | 4.357 | 0.006 | 4.325 | 4.340 | 0.015 | 4.280 | -*^e^* |  |

*^a^* 400MHz, in DMSO- *d_6_*

*^b^* the down - filed chemical shift of the ^1^H NMR spectrum between **PPIX** added with and without lipid.

*^c^* the down - filed chemical shift of the ^1^H NMR spectrum between **PPIX** mixed with 10% of water, and which was added with lipid.

*^d^* the down - filed chemical shift of the ^1^H NMR spectrum between **PPIX** mixed with 16.67% of water, and which was added with lipid of MDCK Cell.

*^e^* the chemical shift was covered by NMR signals of water

**Table S2** Thermodynamic parameters of lipid interaction between **PPIX** and MDCK cells

| Name | K_d_ (µM) | K_a_ (µM^-1^) | ΔH (kJ/mol) | ΔG (kJ/mol) | -TΔS (kJ/mol) |
| --- | --- | --- | --- | --- | --- |
| **PPIX** | 85.1 | 0.01 | -162 | -23.3 | -139 |
